# Development and utilization of new O_2_-independent bioreporters

**DOI:** 10.1101/2023.11.07.566077

**Authors:** Eva Agranier, Pauline Crétin, Aurélie Joublin-Delavat, Léa Veillard, Katia Touahri, François Delavat

**Affiliations:** Nantes Université, CNRS, US2B, UMR6286, F-44000 Nantes, France; Laboratoire Chimie et Biochimie de Molécules Bioactives, Université de Strasbourg/CNRS, UMR7177, F-67081 Strasbourg, France

**Keywords:** bioreporter, Flavin-binding Fluorescent Protein, *Vibrio*, *Rhizobiaceae*, anaerobic condition, anoxia

## Abstract

Fluorescent proteins have revolutionized science since their discovery in 1962. They have enabled imaging experiments to decipher the function of proteins, cells and organisms, as well as gene regulation. GFP and all its derivatives are now standard tools in cell biology, immunology, molecular biology and microbiology laboratories around the world. A common feature of these proteins is their O_2_-dependent maturation allowing fluorescence, which precludes their use in anoxic contexts. In this work, we report the development and *in cellulo* characterization of genetic circuits encoding the O_2_-independent KOFP-7 protein, a flavin-binding fluorescent protein. We have optimized the genetic circuit for high bacterial fluorescence at population and single-cell level, implemented this circuit in various plasmids differing in host range, and quantified their fluorescence under both aerobic and anaerobic conditions. Finally, we showed that KOFP-7 based constructions can be used to produce fluorescing cells of *V. diazotrophicus*, a facultative anaerobe, demonstrating the usefulness of the genetic circuits for various anaerobic bacteria. These genetic circuits can thus be modified at will, both to solve basic and applied research questions, opening a highway to shed light on the obscure anaerobic world.

**Importance:** Fluorescent proteins are used since decades, and have allowed major discoveries in biology in a wide variety of fields, and are used in environmental as well as clinical contexts. GFP and all its derivatives share a common feature: they rely on the presence of O_2_ for protein maturation and fluorescence. This dependency precludes their use in anoxic environments. Here, we constructed a series of genetic circuits allowing production of KOFP-7, an O_2_-independant Flavin-Binding Fluorescent Protein. We demonstrated that *Escherichia coli* cells producing KOFP-7 are fluorescent, both at the population and single-cell levels. Importantly, we showed that, unlike cells producing GFP, cells producing KOFP-7 are fluorescent in anoxia. Finally, we demonstrated that *Vibrio diazotrophicus* NS1, a facultative anaerobe, is fluorescent in the absence of O_2_ when KOFP-7 is produced.

Altogether, the development of new genetic circuits allowing O_2_-independent fluorescence will open new perspective to study anaerobic processes.

## Introduction

Fluorescent proteins have revolutionized our understanding of many biological systems, allowing direct visualization of events on a wide range of spatiotemporal scales. GFP (Green Fluorescent Protein), discovered fortuitously in 1962 (*1*) and its numerous derivatives (RFP, CFP, YFP, eGFP, mTurquoise, mCherry, etc (*2*)) have been particularly used as reporter proteins to address many fundamental and applied questions related to cell visualization, protein-protein interactions, protein localization, biosensor construction, biofilm formation and gene expression dynamics. One of its major advantages relies in the fact that chromophore maturation does not require any cofactor or enzyme, implying that producing the protein *in cellulo* is sufficient to fluorescently tag cells or cell components. However, the main drawback of these proteins is that they require molecular dioxygen (O_2_) for chromophore maturation and thus for fluorescence (*3*). This dependence on O_2_ is a major hindrance to the characterization of systems under low O_2_ tension conditions.

On the other hand, many key microbial metabolic processes occur in hypoxia or anoxia. These can be found in environmental contexts, such as soil (*4*), deep and costal sea waters (*5*) or eutrophic lake waters (*6*), and infectious contexts such as gastrointestinal tracts (*7*) or tumors (*8*). In these O_2_-deprived environments, the use of GFP or derivatives is therefore inefficient. In order to circumvent this pitfall, organic dyes have been experimentally tested, but those generally reveals poorly permeable and/or are cytotoxic, and often accompanied by a high fluorescence background (for a review, see (*9*)). Fluorescent proteins have also been used in anoxia. For example, strict anaerobic bacteria such as *Clostridium difficile* producing the fluorescence protein mCherry can fluoresce when samples are placed in the open air for one hour (*10*). However, this dependence on oxygen can lead to cell lysis in strict anaerobes (*10*), and does not allow its use in real time to monitor the dynamics of biological processes. Other O_2_-independent fluorescent proteins exist, such as UnaG (*11*), which uses bilirubin as a cofactor to fluoresce (*12*). However, they require the addition of this cofactor in the culture medium, making these reporter genes of little interest in complex contexts such as the *in vivo* study of bacteria interacting with its host (symbioses, pathogens). Similarly, *lux* bioluminescence genes, although they can be applied to anaerobes (*13*), do not allow quantification of the expression of genes of interest at the single cell level.

Recently, another class of reporter genes has emerged, and these genes encode Flavin Binding Fluorescent Proteins (FbFPs) (*14, 15*). FbFPs present many advantages: they are small (100-140 amino acids, compared to ∽240 amino acids for GFP), have a fast maturation time and fluoresce at similar wavelengths as GFP (*16*). Moreover, they have the great advantage of not requiring O_2_ to mature and fluoresce. Indeed, these proteins use flavin mononucleotide (FMN), a endogenously-produced cofactor. This O_2_-independent maturation opens the way to the development of new reporter genes for the study of biological systems under anoxic or variable O_2_ tension conditions. Several developed FBFP-derivatives have allowed the study of gene expression of subcellular localization of proteins in different prokaryotic and eukaryotic models (*14, 15*). However, these bioreporter proteins typically exhibit low fluorescence compared to GFP and its derivatives (about 30-fold (*17*)), and this represents a bottleneck for their application.

Recently, a random mutagenesis approach was conducted on a gene encoding the SB2 protein (*14*), an FbFP from *Pseudomonas putida*. After several rounds of enrichment in fluorescent clones, the KOFP-7 protein was obtained (*17*). The *in vitro* purified KOFP-7 protein shows a fluorescence intensity as high as that of GFP, as well as a good stability in acidic media or in the presence of reducing agents. These characteristics make KOFP-7 a good candidate for the development of new reporter genes for application to the study of microbial biological systems under anoxic conditions. However, whether KOFP-7 can be used *in vivo* remains to be demonstrated.

In this study, we present the generation of a series of replicative plasmids, differing in their genetic circuits and bacterial hosts, improved for KOFP-7 production and fluorescence under anaerobic conditions. Moreover, we demonstrated that KOFP-7 based fluorescence can be obtained both in *Escherichia coli* and in *Vibrio diazotrophicus*, demonstrating the versatility of these construction. These genetic circuits will contribute in shedding a new light in previously-dimmed anaerobic microbes, for future basic and applied discoveries.

## Results

### Constructions with KOFP-7 allow in cellulo fluorescence under aerobic conditions

In order to test the possibility of using KOFP-7 as a new reporter protein, we sought to create various plasmids, differing in their RBS region, promoter sequences as well as backbone plasmids, all being designed to constitutively express *kofp-7*. We created 7 plasmids (Fig. 1), all of them being able to replicate in *E. coli*, among which four can replicate in *Vibrionaceae* and *Pseudoalteromonaceae* families (*18, 19*) (pFD115, pFD116, pFD145 and pFD150, the latter being promoterless and served as a negative control), two being able to replicate in *Rhizobiaceae* (pFD141 and pFD148), and one containing the broad host-range origin of replication pBBR1 (*20*) (pFD149).

**Fig. 1.**
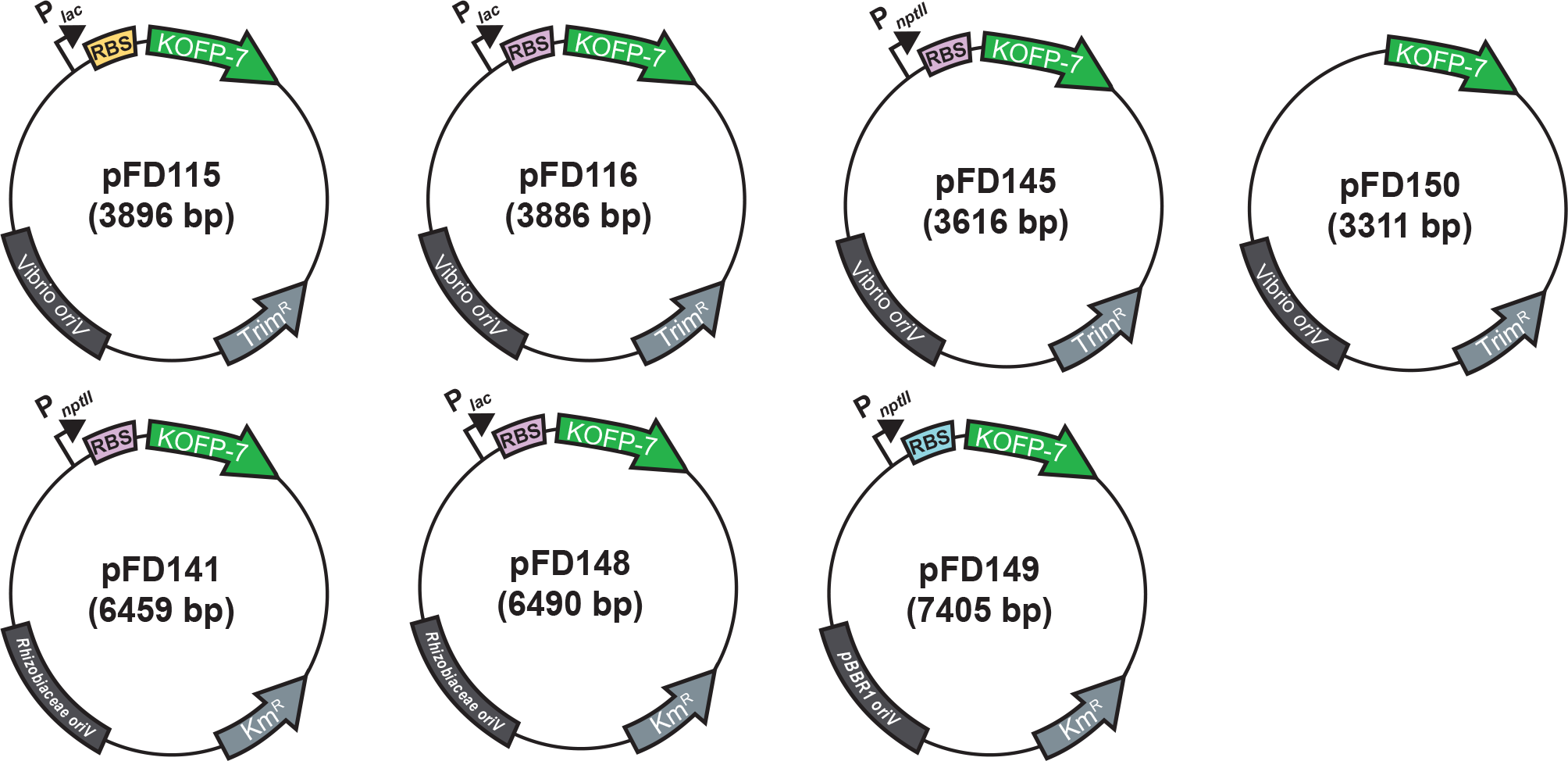
Schematic overview of the constructed plasmids. Identical colors indicate identical sequences. RBS: Ribosome Binding Site. *oriV*: origin of replication.

Our initial screening was performed by quantifying the fluorescence of each of these populations in microplates in the presence of oxygen, all being diluted to the same O.D from overnight cultures. We showed here that while pFD115 fails to produce a fluorescence signal above background, all other constructions allowed fluorescence already at time 0, when cells were in stationary phase (Fig. 2A). Similar conclusions were obtained when the plates were incubated for 5 hours at 37°C before measurement (Fig. 2B), despite all still having a similar final O.D. It is worth noting that pFD115 and pFD116 are very similar, differing only in their RBS region, but the measured fluorescence intensity greatly differs. The highest fluorescence was obtained with the *E. coli* strain carrying pFD145, a plasmid which carries the P_*nptII*_ promoter instead of the P_*lac*_ promoter found in pFD116, the RBS region originally found in pFD086 and the *kofp-7* gene. Interestingly, pFD116 and pFD145 vary only in their promoter sequence, but the measured fluorescence intensity was 1.6 fold higher with the strain containing pFD145. This shows that the P_*nptII*_ promoter is stronger than P_*lac*_ in this genetic context.

**Fig. 2.**
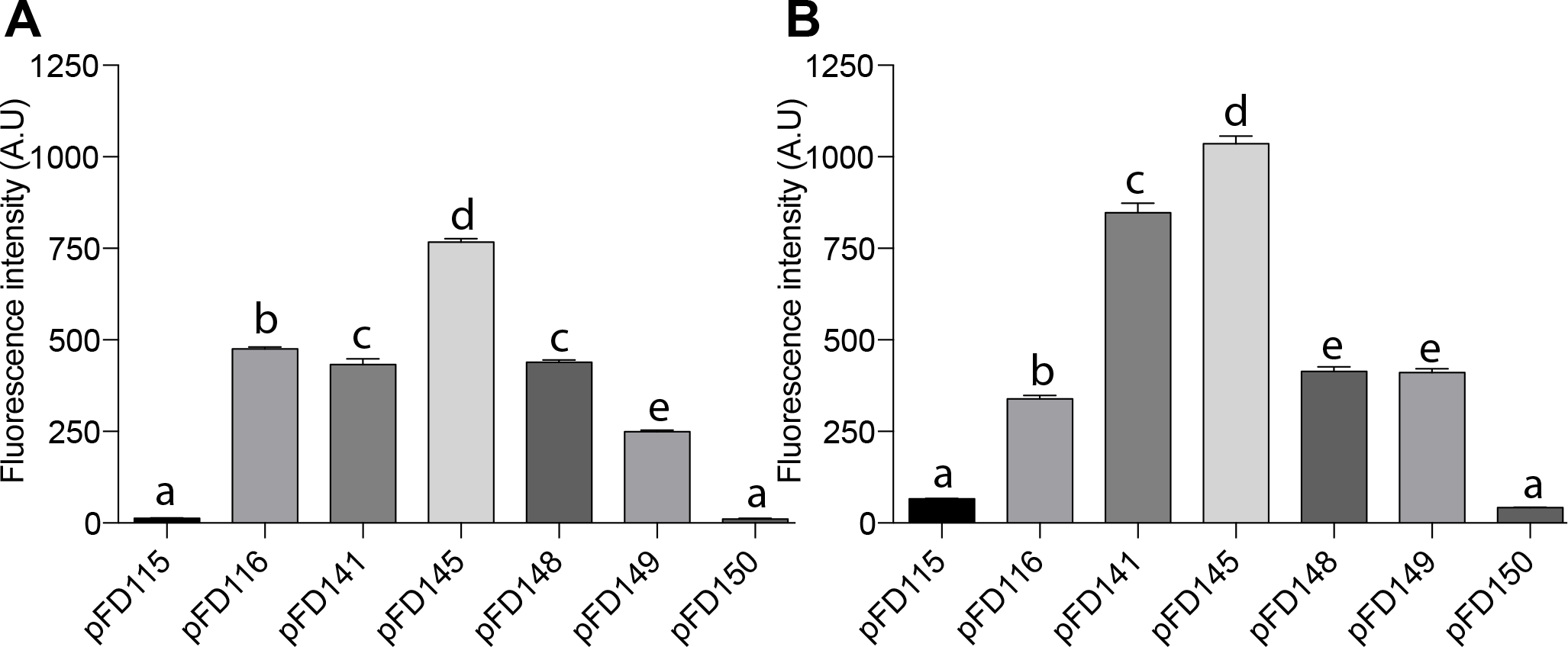
Population-based fluorescence from *kofp-7*-containing strains, measured by fluorimetry after growth under aerobic condition. (A) Fluorescence of *E. coli* carrying the different plasmids measured at time 0. (B) Fluorescence of *E*. coli carrying the different plasmids measured after 5 hours of growth. Presented here are the mean+SEM.

pFD141 and pFD148, both having the same structure and differing only in the inserted promoter (P_*nptII*_ and P_*lac*_, respectively), show intermediate fluorescence intensity, with pFD141 allowing a slightly higher KOFP-7 production (despite being non-significant at time 0). This confirms that P_*nptII*_ is a stronger promoter, also in this genetic context. Finally, the broad-host range vector pFD149, endowed with the P_*nptII*_ promoter shows intermediate results, with the fluorescence being higher than pFD115 and lower than the others. In conclusions, we demonstrate here that the strains containing *kofp-7* as a reporter gene are fluorescent when measured at the population level in microplate. The next step was to quantify the fluorescence level of single cells, using epifluorescence microscopy equipped with a regular FITC-based filter cube.

### KOFP-7-based bioreporters are fluorescent in single bacterial cells

Bioreporters are particularly useful to monitor metabolic processes or to follow the fate of single bacterial cells under various conditions, and epifluorescence microscopy is the corresponding gold-standard. Demonstrating that the newly designed bioreporters can be used at the single cell level was therefore necessary. Cells were grown overnight in LB supplemented with their corresponding antibiotics under aerobic conditions, and fluorescence of single bacterial cells was quantified using in-house Matlab scripts (*21*). Under this condition, single-cell fluorescence measurement are in total accordance with the fluorescence measured from whole populations in microplates (Fig 3A). Specifically, *E. coli* carrying pFD145 performed best, showing the highest per-cell fluorescence intensity, followed by the strain carrying pFD141 and pFD116. Again, *E. coli* cells carrying pFD115 were not fluorescent. This shows that the increase in fluorescence observed in microplate observed for most constructions (Fig. 2) is mostly due to the genetic circuit allowing KOFP-7 production, which differs between plasmids. Under this aerobic conditions, *E. coli* carrying pFD086, differing from pFD116 by the production of EGFP instead of KOFP-7, shows a much higher per-cell fluorescence, as expected from an optimized fluorescent molecule (Fig. 3A). Of note, *E. coli* cells carrying pFD115, pFD116, pFD145 and pFD149 are rod-shaped, as expected from *E. coli* cells (Fig. 3B). However, some cells of strains carrying pFD141 and pFD148 are slightly elongated, possibly reflecting a bacterial stress (Fig 3C). These cells also show intracellular structure resembling inclusion bodies (Fig 3C).

**Fig. 3.**
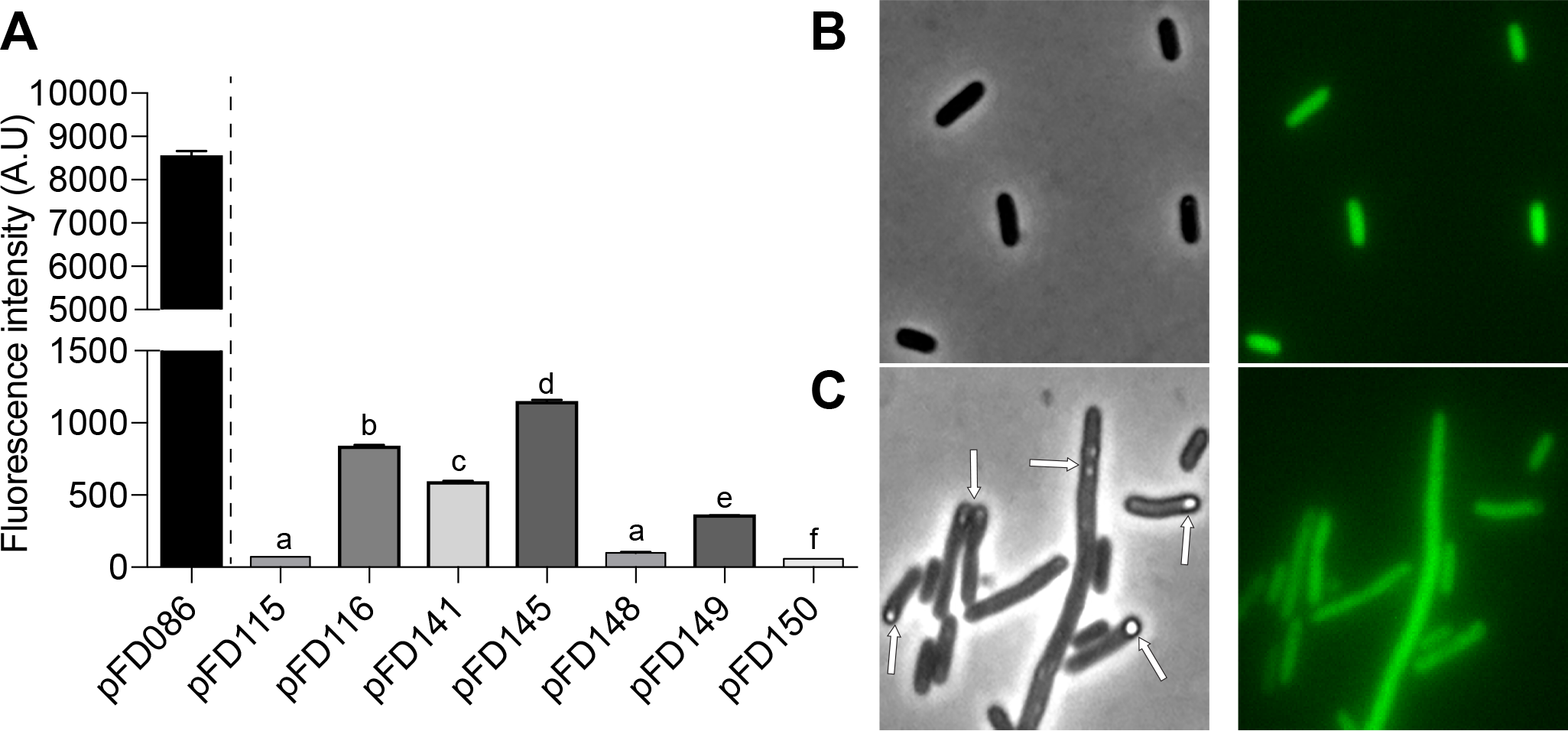
Single-cell-based fluorescence from *kofp-7*-containing strains after growth under aerobic condition. (A) Quantification by epifluorescence microscopy under aerobic condition. Statistical analysis was performed between all *kofp-7*-containing strains. Presented here are the mean+SEM. (B) *E. coli* cells carrying pFD145 and having a classical rod-shape. (C) *E. coli* cells carrying pFD141 and showing aberrant elongated cells, with structures resembling inclusion bodies (see white arrows). Images were scaled to the same brightness.

Finally, we wanted to know the *in cellulo* stability of KOFP-7 fluorescence and its propensity of photobleaching. Therefore, *E. coli* cells carrying pFD145 were deposited on an agarose patch, and cells were exposed to continuous excitation at 482 +/-35 nm. Strikingly, cells experienced rapid photobleaching, with cell fluorescence decreasing sharply within a few seconds, the decrease being visible under the microscope oculars. Photobleaching was measured by quantifying cell fluorescence at time 0 and the fluorescence of the same cells after 5 or 10 seconds of continuous exposure. Cell fluorescence decreased rapidly, reaching a 14-fold decrease after 10 seconds, almost returning to background fluorescence observed from cells carrying the promoterless plasmid pFD150 (Fig 4). Cells carrying pFD086 (and thus producing GFP) exposed to the same treatment were in contrast much less photobleached, with a fluorescence decreasing only 1.2-fold (Fig. 4).

**Fig. 4.**
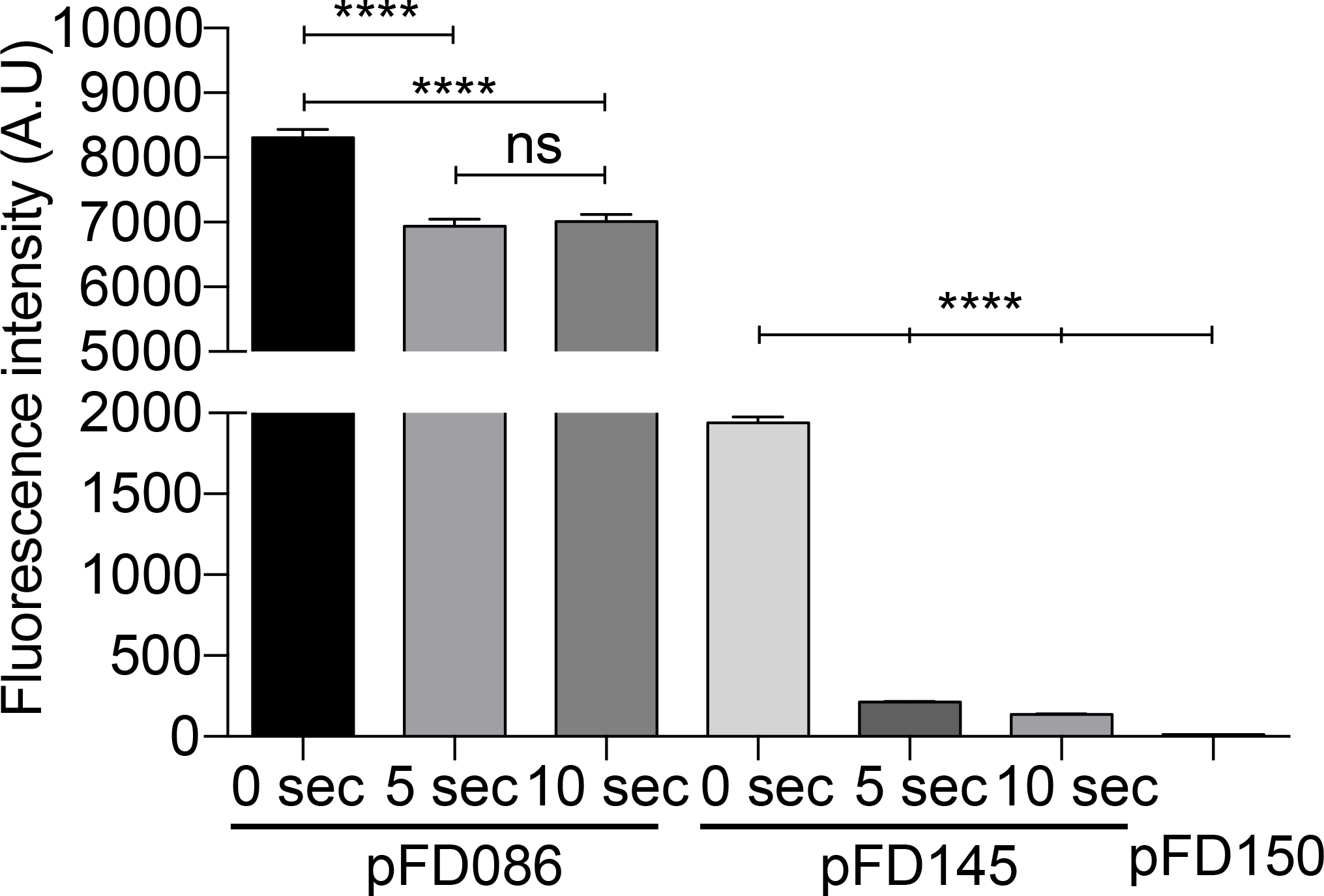
Rapid photobleaching of KOFP-7. Strains producing either the EGFP (pFD086) or KOFP-7 (pFD145) were exposed for 5 to 10 seconds of continuous light, under a microscope equipped with a 482 +/-35 nm filter cube. Fluorescence intensity of single cells was measured at time 0 and after the exposure time. Results from pFD145 were compared to the results of the negative control (strain carrying the promoterless plasmid pFD150). ns: not significant.

### Constructions with KOFP-7 allow in cellulo fluorescence under anaerobic conditions

In order to quantify the fluorescence of the different constructs under anaerobic conditions, LB medium was turned anoxic by flushing the headspace of sealed tubes with N_2_ gas and by the use of the reducing agent thioglycolate. After 24 hours of growth in the absence of oxygen, we quantified the fluorescence from individual cells of the different constructions by epifluorescence microscopy. Under this condition, fluorescence from most constructions was low, with values being close and not significatively different to the negative control (Fig.5A). Exceptions was for the strains carrying pFD116, pFD141, which displays a low-but-significant fluorescence, and the strain carrying pFD145 which showed the highest fluorescence. As expected, the strain carrying a GFP-based plasmid showed a dramatic decrease in fluorescence (compare Fig. 5A with Fig. 3A), as expected from its O_2_-dependancy for GFP maturation. It should be noted that strain growths were very low under this condition, probably because of the absence of electron acceptor for respiration and a very low concentration of fermentative products in the LB medium, hindering the production of energy.

**Fig. 5.**
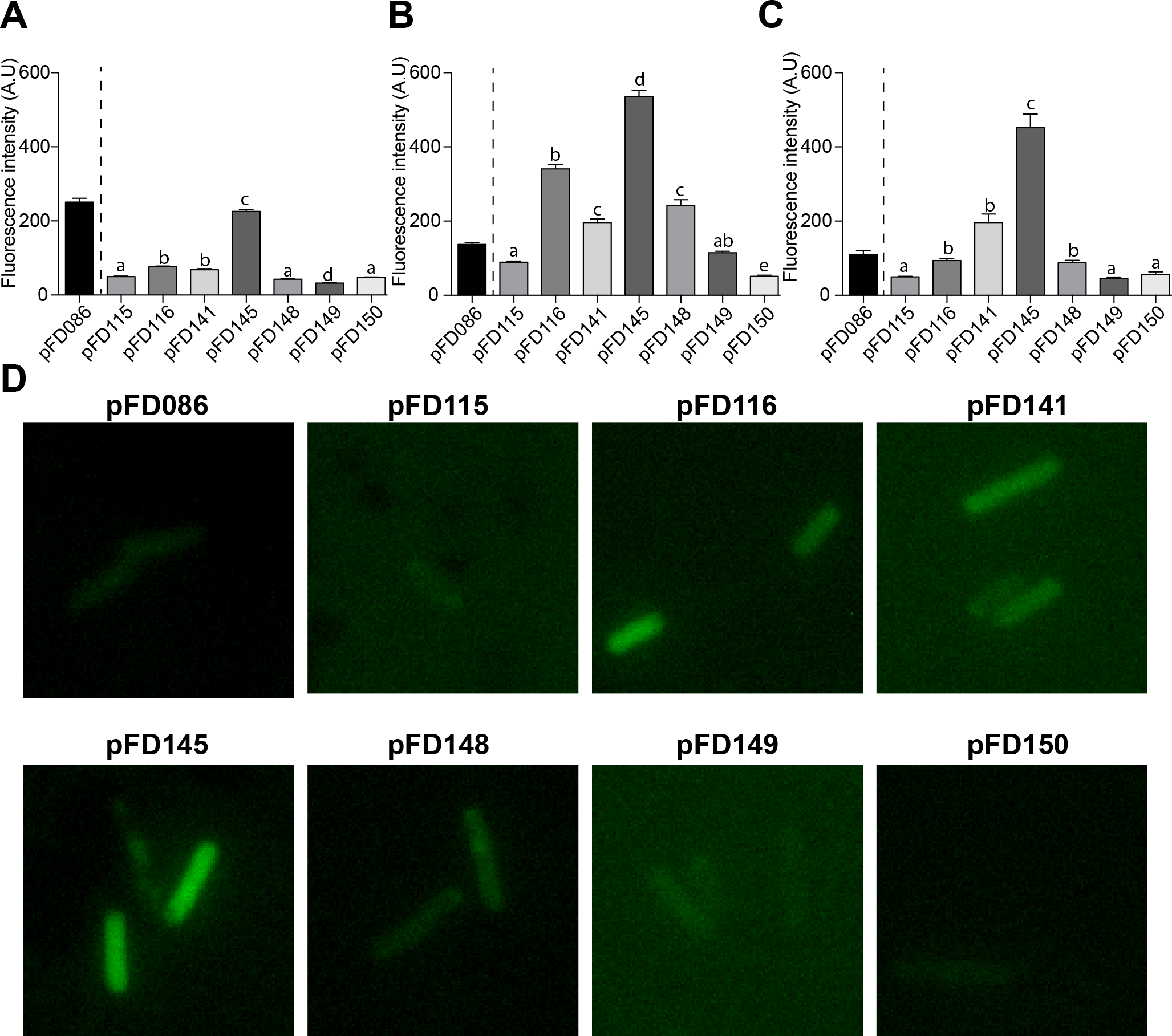
Single-cell-based fluorescence from *kofp-7*-containing strains after growth under anaerobic conditions measured by epifluorescence microscopy. Cells were grown (A) in anaerobic LB conditions ; (B) in anaerobic LB containing NO_3_^-^ as an alternative electron acceptor ; (C) in anaerobic LB containing glucose as a fermentative carbon source. Statistical analyses were performed between all *kofp-7*-containing strains. Presented here are the mean+SEM. (D) illustration of the fluorescence of single cells. Images were scaled to the same brightness.

In a second set of experiments, an alternative electron acceptor (NO_3_ ^-^) was added to anoxic LB medium. Under this condition, fluorescence from KOFP-7 increased significantly, being statistically different from the one of the negative control for all strains, even though the increase in fluorescence due to pFD115 is low (Fig. 5B and 5D). The highest per-cell fluorescence intensity was again obtained for the strain carrying pFD145, followed by the strain carrying pFD116, similar to the results obtained in the presence of oxygen. It should be noted that under this condition, the fluorescence from KOFP-7 is 2.1 fold lower than the one measured in the presence of oxygen (compare Fig. 3A and Fig. 5B). As expected, GFP-containing cells remained dimmed under this condition (Fig 5B and 5D).

In the last set of experiment, every bioreporter was grown without oxygen in LB containing 2% glucose. Fluorescence was similar to the ones obtained in anoxic LB alone for most constructions, with the exception of the strain carrying pFD145, which showed a significant increased fluorescence (Fig. 5C). Under this condition, GFP-containing cells were not fluorescent.

### KOFP-7 based constructions allow cell fluorescence in various bacterial species

Finally, we sought to demonstrate whether KOFP-7 based constructions can be used in other species than *E. coli*. Therefore, we introduced by conjugation the plasmid pFD145, presenting the most promising results in *E. coli*, in *V. diazotrophicus* NS1, a facultative anaerobe with diazotrophic properties. *V. diazotrophicus* cells carrying pFD145 was monitored both in the presence and absence of oxygen in LB medium and we demonstrated that cells are fluorescent, both in the presence (Fig. 6A) and absence (Fig 6B) of oxygen.

**Fig. 6.**
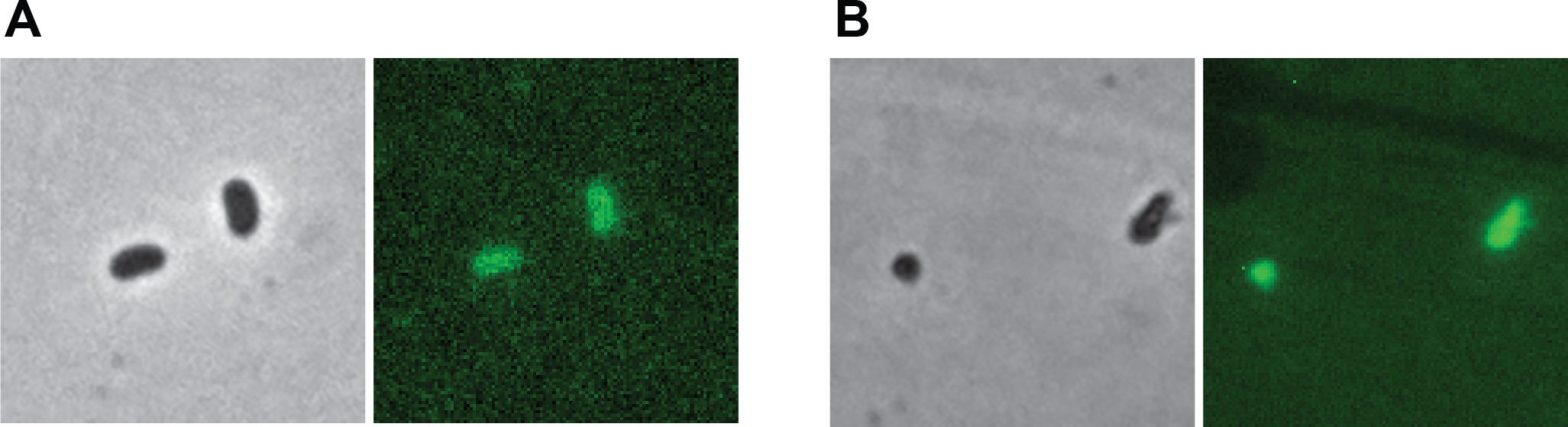
Illustration of the fluorescence of single cells of *V*. *diazotrophicus* containing pFD145 after growth under oxic (A) and anoxic (B) conditions.

## Discussion

Fluorescent proteins have been used since decades to unravel a myriad of biological processes and constitute now a standard working tool in molecular biology, cell biology, microbiology and immunology labs worldwide. The vast majority of these proteins derive from the GFP protein, encoded from the *Aequorea victoria* genome. Many derivatives have been constructed since, differing in their maximum excitation and emission wavelength, their folding speed, their propensity to stay in monomers or their intrinsic brightness (*2*). However, they all have in common their O_2_-dependant maturation, required to turn fluorescent (*3*). This O_2_-dependance represent a major bottleneck in the study of biological processes that take place in the absence of oxygen.

In this study, we present the development of various bacterial bioreporters, which all have in common their O_2_-dependant fluorescence. They rely on the use of KOFP-7, a Flavin-Binding Fluorescent Protein which has been obtained by genetic engineering and which has been reported to present a higher fluorescence and quantum yield, measured *in vitro* from the purified protein (*17*). In order to obtain the highest KOFP-7 production and bacterial fluorescence, various genetic circuits controlling *kofp-7* expression were tested, differing in their promoter strength and RBS regions. We demonstrated here that most of the constructions allowed fluorescence from KOFP-7, both under oxic (Fig. 3) and anoxic conditions (Fig. 5). Moreover, these circuits were implemented into various plasmidic backbones, varying in their origin of replications for *E. coli* (colE1 for pFD141 and 148, R6K for pFD115, pFD116 and pFD145 and pBBR1 for pFD149). These plasmids were also chosen, because they are shuttle vectors and can replicate in bacteria outside *E. coli* : *Vibrionaceae* (*18, 19*) and *Pseudoalteromonaceae* (*18*) for pFD115, pFD116 and pFD145, *Rhizobiaceae* for pFD141 and pFD148, while pFD149 can maintain in various Gram negative bacteria.

Among the tested constructions, the pFD145 plasmid showed the most promising results, allowing the highest population- and individual cell-based fluorescence (Fig 3 and Fig 5). The stability of this fluorescence was therefore tested using this construction. We demonstrated here that KOFP-7 is rapidly photobleached, cells turning almost non-fluorescent after 10 seconds of excitation at 482 +/-35 nm (Fig. 4). This photobleaching can imped the use of KOFP-7 under specific conditions, but most of the applications are performed with an excitation time below the second (*e*.*g*. 100 ms in the present study), decreasing this potential pitfall.

The major advantage of KOFP-7 based bioreporters over GFP-based ones is their capability to fluoresce in the absence of oxygen (*14*). We demonstrated here that the newly constructed bioreporters are indeed fluorescent in the absence of oxygen (Fig. 5). It should be noted that the measured fluorescence is lower than the one obtained with oxygen (Fig 3A). Given the O_2_-independent maturation of the fluorochrome, this decrease in fluorescence is likely due to a decrease in the energetic status of the cells when grown in anaerobic conditions as tested in the present study. Indeed, cells were grown in LB, which is an amino acid-rich medium but otherwise rather poor in saccharides (*22*). In the absence of oxygen or alternative electron acceptor, *E. coli* cells cannot respire and therefore cannot produce ATP by oxidative phosphorylation. In the absence of electron acceptors (Fig. 5A and C), ATP is produced by fermentation. However, amino acids cannot be fermented and the amount of fermentative products is low in LB, limiting this metabolism (*22*). Therefore, the energetic status of *E. coli* when grown anaerobically in LB is low, probably explaining the low cell fluorescence. In line with this hypothesis, the cell fluorescence increased when cells were grown in anoxic condition in LB supplemented with glucose (as a fermentative product, see Fig. 5C) and especially with NO_3_^-^, which served as electron acceptor for anaerobic oxidative phosphorylation. (Fig. 5B). Under the same conditions, GFP-based bioreporters remained dimmed, demonstrating that the increase of KOFP-7-based fluorescence is not due to the addition of oxygen during the injection of NO_3_ ^-^.

Finally, we anticipate that KOFP-7-based constructions can be used to study metabolic and physiological processes in the absence of O_2_. As a proof-of-concept, we monitored the expression of pFD145 in *V. diazotrophicus*, a facultative anaeorbe. We could demonstrate that KOFP7 is produced under anoxic condition and allows cell fluorescence in bacteria other than *E. coli* (Fig. 6). KOFP-7 based constructions are therefore useful tools, functional in various bacterial species and irrespective of the oxygen availability.

To conclude, we present here the development of new set of bioreporters, allowing illumination of bacteria by fluorescence in the absence of oxygen. The developed plasmids can be used in a broad range of Gram negative bacteria and can be used for the study of physiological processes occurring in the absence of oxygen. The current limitation of KOFP-7 is its rapid photobleaching, compensated by its fast maturation time, and its lower intrinsic fluorescence. However, given the recent advances in genetic engineering, derivatives of KOFP-7 or other FbFPs with increased fluorescence will likely emerge in the nearer future, in order to shed light onto the still under-explored world of anoxic environments.

## Material and methods

### Strain and culture conditions

*Escherichia coli* strains and *V. diazotrophicus* NBRC 103148 strains were routinely grown in LB at 37°C and 30°C, respectively. Some experiments using *V. diazotrophicus* were performed in modified MDV medium defined elsewhere (*23*). In order to create anoxic media, 20 ml LB were prepared in 18 cm glass tubes (Dutscher, ref 508232), supplemented with sodium thioglycolate (T0632, Sigma-Aldrich, 10 mg/l final concentration) closed with a sterile blue rubber stopper, and sealed with an aluminum capsule. The head space was flushed for 2 minutes with either argon or nitrogen gas immediately after autoclaving, when the temperature was still at 80°C. If necessary, agar (15 g.l^-1^) or glucose (20 g.l^-1^) were added before autoclaving. Absence of oxygen was controlled by adding resazurin (500 μg.l^-1^ final concentration from a 500 mg.l^-1^ stock solution) in the media before autoclaving. If necessary, trimethoprim (Trim, 10 μg.ml^-1^), kanamycin (Km, 50 μg.ml^-1^), diaminopimelic acid (DAP, 0.3 mM), NaNO_3_ (4,67 g.l^-1^ from an autoclaved 466,7 g.l^-1^ stock solution), 20 g.l^-1^ were added after autoclaving.

For anoxic growth, one colony of each strain was grown overnight in 100ml-erlenmeyer containing 20 ml LB+antibiotic. The next morning, anoxic LB were inoculated with 100 μl of overnight cultures and the antibiotic, using sterile syringes and needles. If necessary 200 μl of NaNO_3_ were added at the same time. Tubes were incubated for 24 hours before fluorescence quantification as described below. The strains, plasmids and primers used in this study can be found in Table S1, S2 and S3, respectively.

### Strain constructions and DNA techniques

Standard procedures were used for all molecular techniques, following the reagent suppliers recommendations. *E. coli* strains were transformed using heat-shock transformation. *V. diazotrophicus* was modified by conjugation, using a recently published protocol (*23*).

### Construction of plasmids with constitutive kofp-7 expression

A synthetic sequence containing the ORF of KOFP-7 was obtained by GeneArt (ThermoFisher Scientific). This sequence contained several attributes, allowing the construction of various plasmids differing in their ribosome binding sites (RBS) and promoter sequences, thanks to restriction sites flanking the to-be-cloned region or by Gibson assembly (using the NEBuilder Hifi DNA assembly master mix, NEB). pFD115 was obtained by replacing the RBS region and *gfp* gene of pFD086 (*24*) with a fragment containing the RBS of pOT1e (*25*) and the *kofp-7* gene, using SphI + BamHI. pFD116 plasmid was obtained by replacing the *gfp* gene with the *kofp-7* gene, using SphI+BamHI. pFD141 was obtained by Gibson Assembly, replacing the *mcherry* gene of pMG103-nptII-mcherry (*26*) with the *kofp-7* gene, using primers 220415-220418. pFD145 was obtained by Gibson assembly, replacing the P_*lac*_ promoter of pFD116 with the P_*nptII*_ promoter of pFD141, using 230214-230217. pFD148 was obtained by replacing the P_*nptII*_ promoter of pFD141 with the P_*lac*_ promoter of pFD116 using XbaI+SphI. pFD149 was obtained inserting the P_*nptII*_ promoter of pFD145 (using XhoI+XbaI) in pFD147 (using SalI+XbaI). pFD150 was obtained by digesting pFD116 and pFD085 with XbaI+PstI, replacing the promoterless *gfp* gene by *kofp-7i*. All constructions were verified by PCR and, when Gibson assembly was done, the absence of mutations, which might have occurred during PCR amplification, has been verified by sequencing. All *in silico* plasmid maps are available upon request.

### Fluorescence quantification by fluorimetry

One colony of each strain containing plasmids allowing constitutive KOFP-7 production was grown overnight in LB+antibiotic. The next morning, OD_600nm_ was measured, 1 ml of each culture was centrifuged, the pellet was resuspended with 1 ml of minimum fluorescent medium (1 l distilled water supplemented with 10 g glucose, 0.1519 g NH_4_Cl, 0.028 g K_2_HPO_4_, 5 g NaCl, 0.5 g yeast extract and 0.1 g tryptone (modified from (*27*)). The pH was adjusted to 7 with a solution of HCl (0.2 M) or NaOH (0.2 M), which displays a lower autofluorescence than LB (*27*), and further diluted with the same medium to reach OD 0.1. 200 μl of these OD-adjusted cultures were inoculated in 96-well black polystyrene plates (ref 655900, Greiner Bio-One) in quadruplicates and fluorescence was measured using the TECAN Infinite M1000, set with an excitation wavelength of 450 nm and a emission wavelength of 496 nm. Wells filled with minimum fluorescent medium + antibiotic without bacteria were used as a control to remove the fluorescence background of the medium. Microplates were incubated at 37°C with shaking, and fluorescence was measured every hour over 5 hours.

Every experiment was done in biological triplicates (3 independent days), with 4 technical replicates per biological replicate, every biological replicate giving a similar result. Presented here are the results obtained from one biological replicate.

### Fluorescence quantification by microscopy

Cultures were grown overnight under aerobic conditions or for 24 hours in anoxia. Aliquots of these cultures were subsequently used for fluorescence microscopy. Briefly, agarose patches were prepared by melting 1% agarose in water, followed by incubation at 50°C to equilibrate the temperature. 600 μl of molten agarose were deposited onto a microscopy slide, and a second microscopy slide was immediately added on top of the liquid. 10 minutes later, the patch was recovered by sliding the 2 microscopy slides and drops of the overnight culture were deposited on the agarose patch before being covered with a coverslip. Pictures were acquired from the overnight cultures using an Eclipse Ni-E microscope (Nikon) equipped with a 100X Apochromat oil objective, a FITC-3540C filter cube (482 +/-35 nm excitation) at 100 ms exposure time. Fluorescence of single cells was quantified using an in-house written Matlab script (*21*) from at least 8 independent pictures. Every condition was tested at least in duplicates, with similar results. Presented here are results from one replicate.

In order to observe KOFP-7 based fluorescence in *V. diazotrophicus*, we grew cells *V. diazotrophicus* in LB+Trim. 100 μl of these overnight precultures were used to inoculate 20 ml LB+Trim either under aerobic condition (in 100 ml-erlenmeyer with shaking) or under anaerobic condition (18 cm glass tubes as described above) and 200 µl of NaNO_3_^-^. After 12 hours (for aerobic cultures) or 24 hours (for anoxic cultures), drops of the cultures were deposited onto agarose patch and immediately monitored by epifluorescence microscopy.

### Statistical analyses

All statistical analyses were performed using GraphPad Prism; *p* values <0.05 were considered statistically significant.

## Acknowledgments

This work was supported by the “Rising Star” (Pays de la Loire Region) and “DBM” (CNRS-INSB) grants, awarded to François Delavat. Pauline Crétin was supported by the CNRS-MITI (GdR OMER). Authors would like to thank Jordan Vacheron (DMF, University of Lausanne) and Eric Giraud (IRD Montpellier) for sharing the pOT1e and pMG103-nptII-mcherry plasmids, respectively.

## Author contributions

François Delavat: Conceptualization, Data curation, Formal analysis, Funding acquisition, Investigation, Methodology, Project administration, Resources, Validation, Visualization, Writing – original draft | Eva Agranier, Pauline Crétin, Aurélie Delavat, Katia Touahri: Data curation, Formal analysis, Methodology, Investigation, Writing – review and editing | Léa Veillard: Formal analysis, Writing – review and editing

